# An Engineered T7 RNA Polymerase for efficient co-transcriptional capping with reduced dsRNA byproducts in mRNA synthesis

**DOI:** 10.1101/2022.09.01.506264

**Authors:** Mathew Miller, Oscar Alvizo, Chinping Chng, Stephan Jenne, Melissa Mayo, Arpan Mukherjee, Stuart Sundseth, Avinash Chintala, Jonathan Penfield, James Riggins, Xiyun Zhang, Antoinette Sero, Justin Dassie, Neil Leatherbury, Scott Baskerville, Gjalt Huisman

## Abstract

Messenger RNA (mRNA) therapies have recently gained tremendous traction with the approval of mRNA vaccines for the prevention of SARS-CoV-2 infection. However, manufacturing challenges have complicated large scale mRNA production, which is necessary for the clinical viability of these therapies. Not only can the incorporation of the required 5’ 7-methylguanosine cap analog be inefficient and costly, in vitro transcription (IVT) using wild-type T7 RNA polymerase generates undesirable double-stranded RNA (dsRNA) byproducts that elicit adverse host immune responses and are difficult to remove at large scale. To overcome these challenges, we have engineered a novel RNA polymerase, T7-68, that co-transcriptionally incorporates both di- and tri-nucleotide cap analogs with high efficiency, even at reduced cap analog concentrations. We also demonstrate that IVT products generated with T7-68 have reduced dsRNA content.

---

There have been many advancements in mRNA therapeutics in recent years, with significant improvements in protein translation, modulation of immune responses, formulation, and delivery. As a result, mRNA therapeutics are clinically progressing in, immuno-oncology, and prophylactic vaccines [1]. The rapid development and overall efficacy of the [BNT162b2](https://clinicaltrials.gov/ct2/show/NCT04380701) and mRNA-1273 SARS-CoV-2 vaccines, as well as their potential to be quickly and flexibly deployed in response to emerging variants, has established mRNA as a critical molecule for global health [2]. At the same time, recent history has highlighted the costs and challenges of producing mRNA at large scale, limiting the worldwide accessibility of mRNA therapeutics [1]. Importantly, mRNA capping and downstream processing of the mRNA to remove immunogenic double stranded RNA (dsRNA) contribute significantly to the overall cost and complexity of production.

mRNA capping occurs in eukaryotes at the 5’ end of a nascent transcript with the addition of a 7-methylguanosine (m7G) cap structure through a 5’-5’ triphosphate linkage by a cap-synthesizing complex associated with RNA polymerase II [3]. The cap is critical for mRNA metabolism in the cell: It recruits ribosomes to the 5’ end of the mRNA to initiate ribosome scanning and translation of the encoded protein. It protects mRNAs from 5’ exonucleases to regulate RNA half-lives *in vivo*, and it prevents triggering an antiviral response through adjacent 3’-O methylation residues that contribute to self-RNA recognition [3]. To ensure their clinical safety and efficacy, mRNA therapeutics must also be generated with a cap structure. In fact, the RNA capping efficiency is a critical quality attribute (CQA) in the manufacture of mRNA at scale [4].

Synthetic mRNAs are produced through in vitro transcription (IVT) from a DNA template. Here, the RNA polymerase from bacteriophage T7 (T7RNAP) is widely used due to its robust yield and high processivity. In the co-transcriptional capping process, a di- or tri-nucleotide m7G cap analog is added to the IVT reaction. The cap analog base-pairs with the template at the transcription initiation site and is incorporated at the 5’ end of the nascent transcript. However, GTP present in the IVT nucleotide pool competes with the cap analog for incorporation at the +1 position, because wild-type (WT) T7RNAP has no selectivity for dinucleotide cap analogs over GTP. To overcome this, a 4-fold molar excess of the cap analog over GTP is required to bias the reaction towards initiation with the cap analog and reach ∼70% capping efficiency [5]. This high cap analog concentration, coupled with the low GTP concentration increases process cost and decreases final mRNA yield, respectively. An alternative process, often referred to as enzymatic capping, uses the vaccinia capping enzyme (VCE) complex to add a cap structure to mRNAs post-IVT. However, this process does not always reach completion [6] [7]. As it uses an additional enzyme with unique reaction condition tolerances, it also requires an mRNA isolation step between the IVT and capping reactions, resulting in yield losses, increased complexity and ultimately expense.

Although WT T7RNAP demonstrates robust mRNA yield generation during IVT, it produces unwanted double-stranded RNA (dsRNA) byproducts through its templated RNA-dependent RNA polymerase activity [8]. dsRNA byproducts stimulate host immune responses by activating pattern recognition receptors (PRRs), such as endosomal-bound Toll-like receptor-3 (TLR3), and the cytosol localized Retinoic acid-inducible gene I (RIG-I) and melanoma differentiation-associated protein 5 (MDA-5). Additionally, dsRNA activates RNA-dependent protein kinase (PKR), leading to the phosphorylation of α-subunit of translation initiation factor-2 (eIF-2α), thereby inhibiting translation [9]. These off-target immune responses reduce the safety and efficacy of mRNA therapies [10] [11]. In mRNA production, downstream chromatography steps are required to remove contaminating dsRNA products from the desired transcript. HPLC purification is the gold standard for small-scale mRNA production, as it removes IVT contaminants such as abortive transcripts and dsRNA [12]. However, this method does not scale for mRNA mass production [13]. Other methods for removing dsRNA or reducing its generation have recently been described. These include cellulose chromatography dsRNA removal [14], sequence engineering by Uridine-depletion of the coding sequence (CDS) [5], and high temperature IVT using a thermostable T7RNAP [15]. However, each of these strategies may not be sufficient in all cases, and a more universal solution that does not require significant process change is still needed.

To this end, we have engineered a novel RNA polymerase with selective incorporation of cap analogs to address the challenges of efficient co-transcriptional capping, resulting in a 4-fold reduction in the required cap concentration during IVT. In parallel, dsRNA byproduct generation in the evolved variant was decreased more than 50-fold, translating into reduced immunogenicity and increased expression in cell-based reporter assays.

## RESULTS

### mRNA capping efficiency for dinucleotide analogs

The commercially available polymerase T7-68 is a directed evolution variant derived from wild-type T7 RNA polymerase. This variant was selected during HTP screening of directed evolution libraries for increased incorporation of dinucleotide cap analogs, while maintaining mRNA yield.

Two dinucleotide cap analogs were selected for capping evaluation studies using T7-68 and WT T7 RNA polymerases. Unlike m^7^GpppG and other dinucleotide analogs, the anti-reverse cap analog m_2_^7,3’-O^GpppG (ARCA) (**S1**) incorporates in only the forward orientation, by virtue of the 3’-O methylation on the methylated guanosine. This analog is widely used because higher overall capping efficiencies are achieved because of this directional incorporation [16]. The cap analog m^7^Gpppm^7^G (sCap) (**S2**) is symmetrical around the 5’-5’ triphosphate linkage, so that incorporation in either orientation provides a 5’ 7-methylguanosine Cap-0 structure [17]. Like ARCA, when sCap is incorporated into the mRNA during initiation, its orientation is correct in 100% of instances. The sCap analog is incorporated with a capping efficiency equivalent to other cap analogs, including ARCA derivatives, while supporting high translation efficiency relative to other analogs [17].

Two reporter constructs, a GlmS ribozyme and a firefly luciferase mRNA, were used to evaluate capping efficiency. mRNA encoding the GlmS ribozyme serves as a convenient reporter, self-cleaving to generate a small 16-mer 5’ analyte upon induction with the ligand glucosamine-6-phosphate (Fig 1A). Cleavage of the reporter depends on sequences internal to the ribozyme structure and immediately adjacent to the cleavage site and is not expected to be influenced by capping at the 5’ end of the mRNA. The capped (m7Gppp-) and uncapped (ppp-) species were resolved using denaturing polyacrylamide gel electrophoresis and visualized by GelGreen staining.

**Figure 1:**
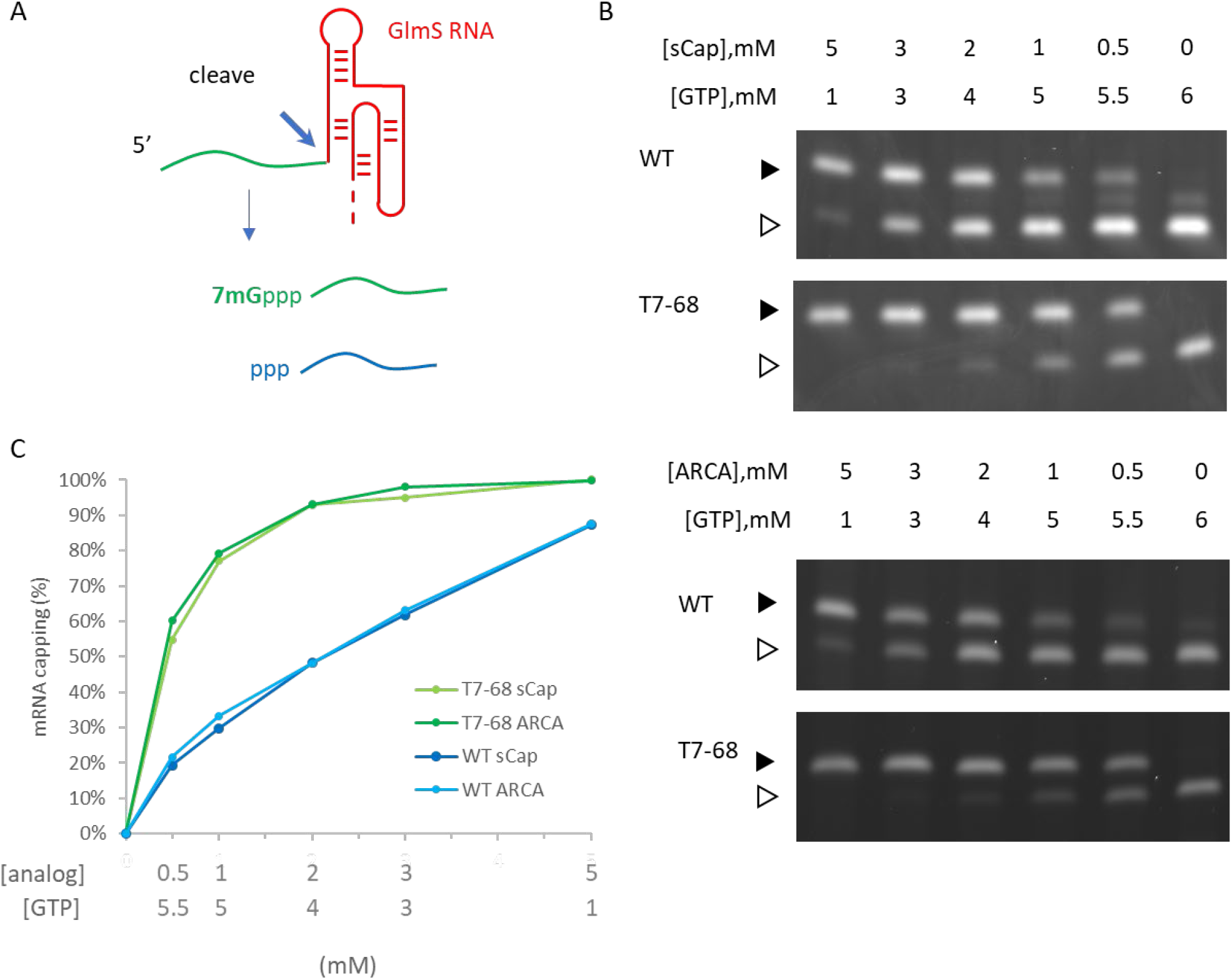
Dinucleotide capping efficiency comparison for sCap and ARCA on a model riboswitch transcript. The WT and T7-68 RNA polymerases were used to transcribe the riboswitch reporter **(A)** using sCap or ARCA analogs. **(B)** The cap analogs and GTP were titrated between 0.5-5mM. Capped (closed triangle) and uncapped (open triangle) 5’ cleavage fragments were resolved on a 15% PAGE-Urea gel. **(C)** Densitometry was used to integrate capped and uncapped species, and % mRNA capping for the WT and T7-68 polymerases is graphed for the full titration.

Co-transcriptional capping IVT reactions using dinucleotide analogs are often assembled with a high ratio of cap:GTP to ensure efficient incorporation of the cap analog using the non-selective WT RNA polymerase. We performed a titration of both the ARCA and sCap analogs from 0-5 mM, with the total concentration of cap and GTP in each reaction equal to 6mM, to generate a range of Cap: GTP ratios in the series. Bands representing the capped and uncapped species were well-resolved for the T7-68 polymerase (Fig 1B). An additional species with an intermediate mobility was observed for the WT T7RNAP, which may represent an additional nucleotide incorporation “stutter” event during transcription initiation. This band was present in the no cap condition, and it is inversely correlated with the cap analog concentration. Interestingly, this intermediate band was not observed in T7-68 reactions run under the same conditions.

Quantitative analysis was performed by gel densitometry (Fig 1C). Across the broad range of Cap analog: GTP ratios tested, the ARCA and sCap analogs produced nearly identical capping efficiencies. For both analogs, the capping efficiency was significantly higher for T7-68 relative to the WT polymerase at a given Cap:GTP ratio. At the highest titration with 5mM Cap and 1mM GTP, the capping efficiency was ∼100% (uncapped was undetectable) for T7-68, while the WT polymerase achieved 88% capping. The greatest improvements in capping efficiencies were observed at low Cap:GTP ratios.

We next evaluated capping efficiency on a 1.8kb firefly luciferase mRNA with a length more representative of therapeutic mRNAs. In this assay, an orthogonal cleavage method was used: an engineered DNA enzyme was used to site-specifically cleave the mRNA, releasing 13-mer analytes for analysis by denaturing Urea-PAGE. As before, we assembled a titration of both the ARCA and sCap analogs from 0-5 mM, with the total concentration of Cap and GTP in each reaction equal to 6mM. Bands corresponding to the capped and uncapped species resolved well, with no evidence of a significant intermediate “stutter” product in either the WT or T7-68 RNA polymerases, at any cap concentration (Fig 2A and 2B).

**Figure 2:**
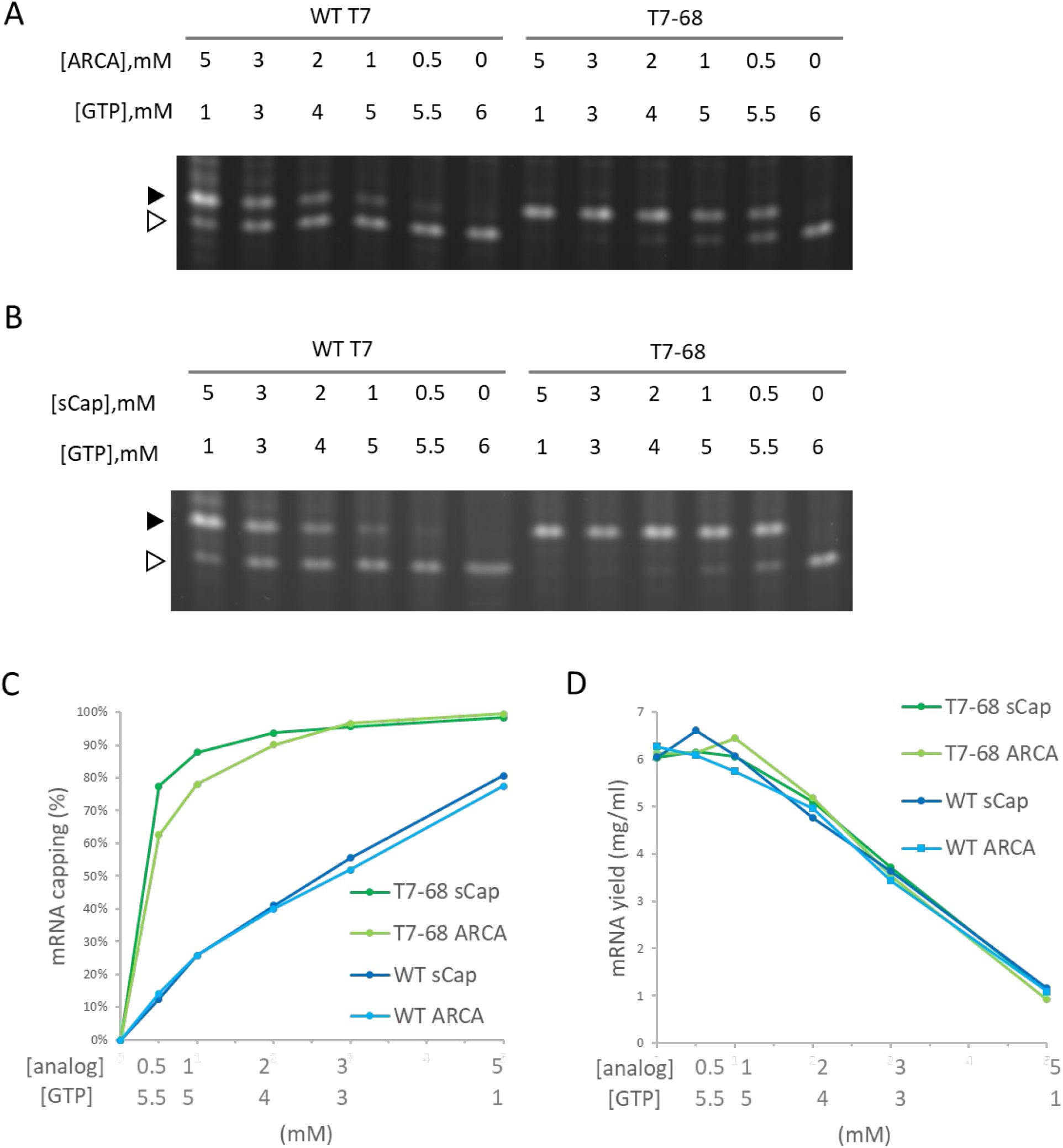
Dinucleotide capping efficiency comparison for sCap and ARCA on a Luciferase reporter mRNA. transcript. The WT and T7-68 RNA polymerases were used to transcribe a luciferase mRNA reporter using cap analogs titrated between 0.5-5mM, with the total of the cap analog + GTP at 6mM. (A) sCap and (B) ARCA analogs were cleaved using a DNAzyme and visualized on 15% PAGE gels. Capped (closed triangle) and uncapped (open triangle) 5’ fragments are labeled. (C) Capped and uncapped fragments were integrated to calculate % capping efficiency for each point in the titration (D) mRNA yields for each point in the titrations were measured by the Quant-iT RNA Broad range assay

Quantitative capping efficiency was similar, though not identical, between the two cap analogs, with slightly higher sCap efficiencies observed at lower Cap:GTP ratios (Fig 2C). Capping efficiency with the sCap analog was 99% for T7-68, compared to 81% for the WT polymerase at the 5mM Cap: 1mM GTP ratio. At an intermediate cap ratio of 1mM Cap: 5mM even greater differences in capping were observed with 88% for T7-68 and 26% for WT T7 RNAP.

We quantified mRNA yield to evaluate the tradeoff between mRNA capping efficiencies and yield as Cap:GTP ratios are varied in co-transcriptional capping batch reactions. mRNA yield for each reaction was determined on quenched IVT reactions using a fluorescent intercalating dye assay. The yield for all four reactions was very similar across all conditions, and roughly correlated with the concentration of GTP in the reaction (Fig 2D). Notably, T7-68 allowed reactions with capping at 88% or higher to proceed with almost no loss in IVT yield (∼ 6 mg/ml), whereas WT T7 RNAP achieved only 77-81% capping under a condition which significantly decreased yield (1mg/ml, 5mM sCap: 1mM GTP). The T7-68 polymerase allowed significant flexibility for optimizing capping efficiency with total mRNA yields > 4mg/ml in a batch reaction.

### Capping efficiency with a trinucleotide analog

The trinucleotide cap analog CleanCap AG **(S1)** improves capping efficiency over dinucleotide analogs because it is selectively incorporated by WT T7RNAP over native nucleotides during initiation. This may be due to the second base-pairing interaction between the cap analog and DNA template during transcription initiation [5]. To determine whether T7-68 further improves capping efficiency with this trinucleotide analog, IVT reactions including 0.5 to 4 mM CleanCap AG were performed using an appropriate firefly luciferase reporter using an “AGG” initiator sequence downstream of the T7 promoter. Because CleanCap AG reactions do not require GTP starvation to achieve high capping efficiency, GTP was held constant at 5mM.

As for the earlier described GlmS riboswitch reporter, an intermediate MW band was observed, which may result from a polymerase “stutter” during initiation of uncapped RNAs. This stutter is present in the no cap condition and absent at higher cap loadings. It is more prominent for the WT polymerase, but also detectable for the T7-68 polymerase in the no-cap control (Fig 3A).

**Figure 3:**
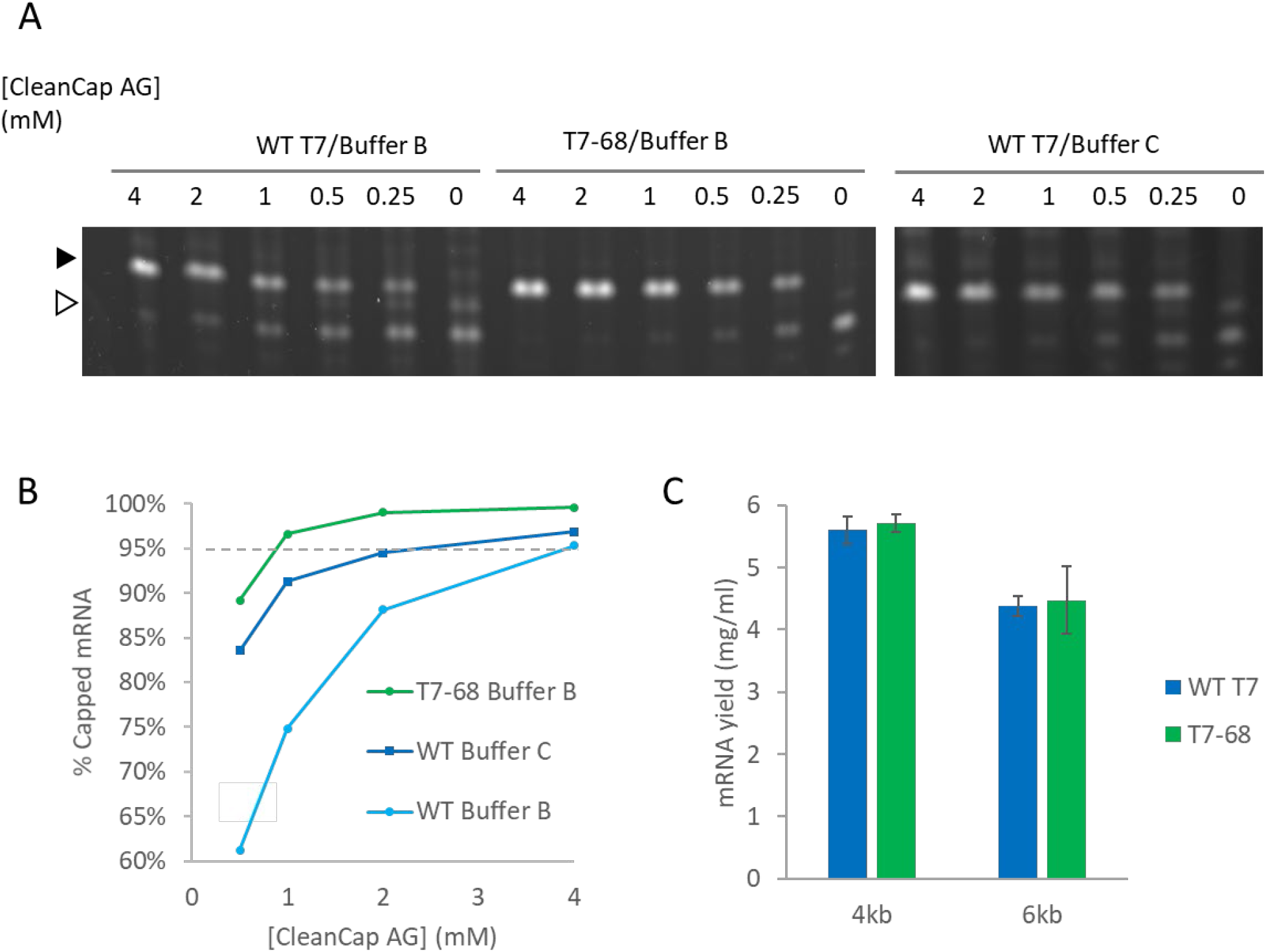
Trinucleotide analog capping efficiency using CleanCap AG with a luciferase mRNA. The WT and T7-68 RNA polymerases were used to transcribe a luciferase reporter using Cleancap AG (0-4mM), and buffers indicated. (A) Capped (closed triangle) and uncapped (open triangle 5’ cleavage DNAzyme cleavage products were resolved on a 15% PAGE gel (B) Gel densitometry was used to quantify mRNA capping efficiency, plotted for each cap concentration. (C) IVT yield was compared for the WT and T7-68 polymerases on 4kb and 8kb templates using 5mM N1mψ.

Using the recommended conditions for co-transcriptional capping with WT T7 RNA polymerase and CleanCap AG (Buffer C, 4mM CleanCap AG), we observed 97% capping efficiency on the luciferase mRNA (Fig 4B), consistent with the 94% value reported from the literature [18]. Maximizing capping efficiency to ∼95% or greater is important to avoid the RIG-I mediated response to uncapped 5’ triphosphate RNA [19]. Notably, the capping efficiency for the WT polymerase is reduced to below 95% at reduced concentrations of CleanCap AG. In its optimized buffer (BufferB), T7-68 achieved 97% capping with 1mM CleanCap AG, equivalent to the efficiency for the WT polymerase with 4mM Cap, for a 4-fold improvement in cap loading (Fig 4B). When T7-68 was loaded at 4mM CleanCap AG, no uncapped products were detectable, providing an opportunity for further improvements in capping efficiency. When compared in the same reaction buffer (Buffer B), even larger differences were observed between the WT and T7-68 polymerase.

**Figure 4.**
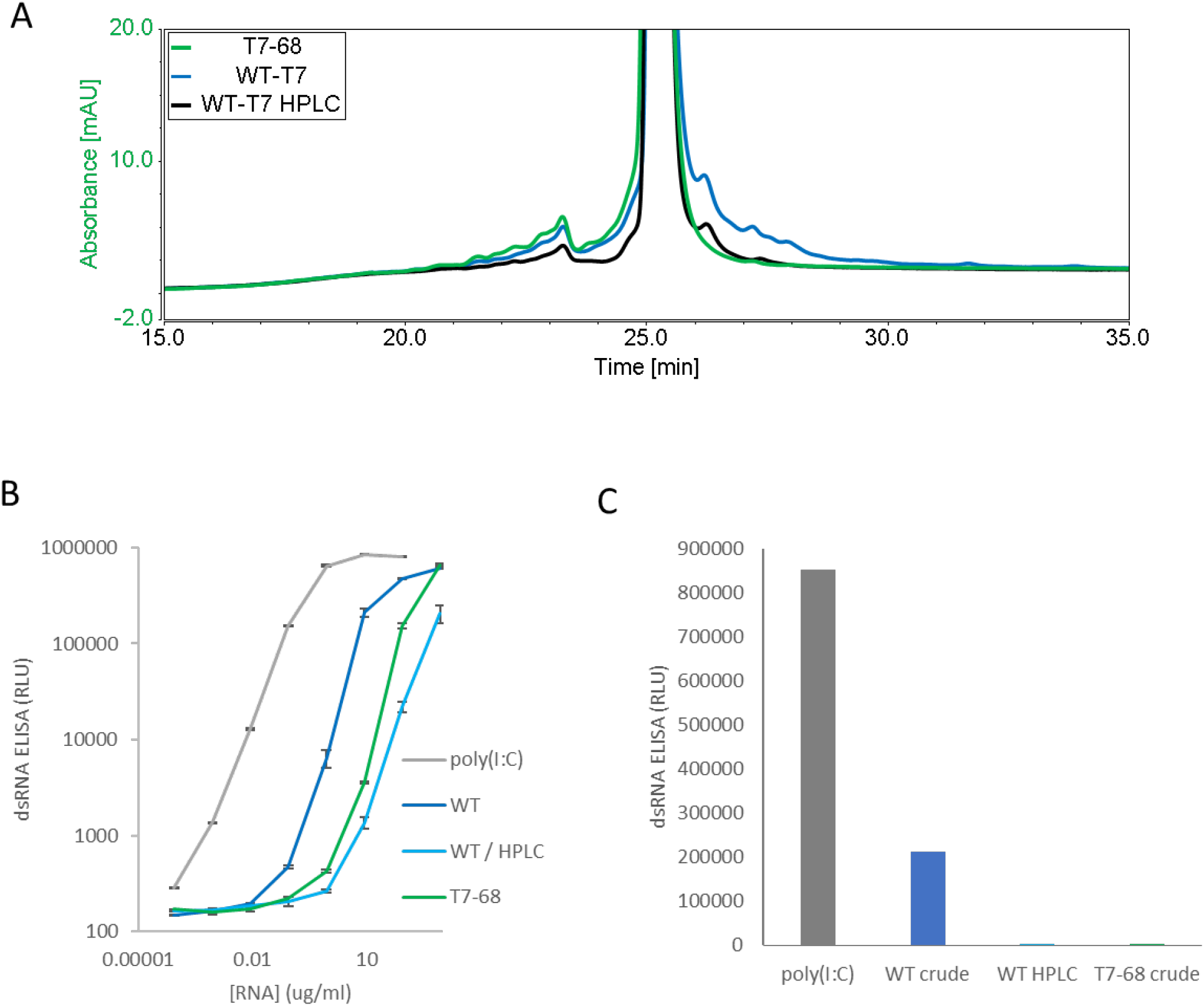
Double stranded RNA produced by T7-68 and WT T7 RNA polymerases. (A) Analytical HPLC chromatography of LiCl precipitated (Crude) or HPLC-purified mRNAs. Traces are normalized by total intensity to allow comparison of 3’ extension products with longer retention times. (B) A dsRNA-specific sandwich ELISA was used to analyze RNAs in panel A for total dsRNA content, using a titration series (log/log plot) (C) A linear plot of dsRNA ELISA signal for mRNA samples at the loading of 9 ng/ul, for which the greatest separation of signal was observed between purified samples and poly I:C control.

### mRNA yield with modified nucleotides

N-1-methylpseudouridine (N-1mψ) is a uracil analog commonly incorporated into mRNA therapies to abrogate the immune-response to unmodified mRNAs [20]. In its optimized buffer, T7-68 produced similar yields relative to the WT polymerase run in a standard reaction buffer (NEB) when tested on 4kb and 6kb templates with 5mM N-1mψ substituted for UTP (Fig 4C). Thus, T7-68 has maintained the ability to utilize the deimmunizing nucleotide over a broad range of IVT template lengths (4-6kb), under a process condition which achieves efficient capping using a reduced input of the cap analog, without sacrificing IVT yield.

### Transcription fidelity

Therapeutic mRNAs must be produced with minimal error to ensure translation of the expected protein. To assay transcription fidelity, we employed an unbiased, direct sequencing approach (Fig S2). WT and T7-68 polymerases were used to transcribe a luciferase template with 0.5 mM sCap. cDNAs were cloned, sequenced, and analyzed to measure RNA polymerase fidelity. The observed error rate for T7-68 (8.2×10^−5^) was statistically indistinguishable from the WT polymerase (8.1×10^−5^), (p>0.05) **(Table 1)**, and was consistent with the reported error rate of 5×10^−5^ for WT T7RNAP [21]. Other T7 polymerase variants with increased error rates (T7-79) were observed in the assay, and these were not selected for further characterization.

**Table 1.** Polymerase Error Rates were measured for T7 RNA Polymerase and derived variants by sequencing cDNA clones. Indels and SNPS were counted in transcribed regions of a luciferase reporter gene, and mutation rates were calculated using to the total number of bases observed for sequenced cDNA clones. A one-tailed binomial test was applied to the mutation rates observed to evaluate statistical significance observed error rate deviations for variant polymerases from the observed WT error rate.

| Polymerase | total bases<br>sequenced | indels | SNPS | total errors | Mutation rate | Binomial test<br>(vs WT value) |
| --- | --- | --- | --- | --- | --- | --- |
| T7-WT | 197472 | 1 | 15 | 16 | 8.1E-05 | 0.533 |
| T7-79 | 195840 | 0 | 34 | 34 | 1.7E-04 | **0.00005** |
| T7-68 | 218688 | 0 | 18 | 18 | 8.2E-05 | 0.505 |

### dsRNA contaminants from IVT

In addition to the desired full-length mRNA, WT T7RNAP generates double stranded RNA (dsRNA) byproducts through two general mechanisms: Antisense RNA products formed during IVT hybridize to the sense mRNA, creating dsRNA duplexes [22], and 3’ extension products longer than the expected runoff transcript are produced when the 3’ end of an mRNA self-hybridizes in *cis*, and T7RNAP extends via an intrinsic RNA-dependent RNA-polymerase activity to form regions of self-complementary dsRNA [23] [24]. Because dsRNA byproducts are known to stimulate an adverse immune response, which concomitantly reduces translation efficiency [25], we characterized the dsRNA byproducts produced by the WT and T7-68 polymerases.

HPLC is an established analytical method to identify 3’ extension products, as well as a small-scale preparative method to isolate the expected mRNA product from dsRNA contaminants [12]. mRNAs were transcribed from a 1.4kb mRNA using N1mψ, and either the WT T7 or T7-68 polymerase with CleanCap AG at 5mM and 2mM, respectively. Capping for both samples was measured at 99% via quantitative LC-MS analysis (TriLink). Higher molecular weight 3’ extension products were observed in the WT IVT sample, and these were significantly reduced in the HPLC-purified WT sample, with two small peaks remaining after purification (Fig 4A). In contrast to the WT samples, the crude T7-68 sample had the lowest amount of high MW RNA, and lacked distinct peaks with longer retention times than the expected mRNA (Figure 4A, S3). Baseline separation between the full-length mRNA and 3’ extensions was not achieved, so these integrations likely underestimate the reductions in 3’ extensions for the crude T7-68 and WT-HPLC samples.

Total dsRNA content was assayed by sandwich ELISA assay, using the dsRNA-specific K1 and K2 antibodies [26]. This assay detects both antisense and 3’ extension dsRNAs. mRNA produced by T7-68 generated 59-fold less dsRNA signal than mRNA produced from WT T7 in the linear range of the assay(Fig 4B and 4C). A similar reduction was observed from HPLC purification of the WT T7 produced mRNA (Fig. 4B, 4C). A second 4kb reporter transcript was assayed for dsRNA in a qualitative dot blot ELISA assay using the J2 dsRNA-specific antibody [26], and lower dsRNA was observed in crude mRNAs transcribed by T7-68 relative to the WT polymerase (Fig S4). Because the same mRNAs were transcribed by both polymerases under the same buffer and reaction conditions, the reduction in dsRNA signal in the T7-68 reaction is polymerase-intrinsic. Taken together, the HPLC and dsRNA ELISA results indicate a significant reduction in dsRNA production using the T7-68 variant.

### Immune stimulation by dsRNA contaminants

We next assessed immune stimulation by synthetic mRNAs in cell-based assays. RIG-I monitors the 5’ end of mRNAs in the cytosol and nucleus, and binds selectively to viral RNA features shared with synthetic mRNAs. Such features include RNAs with uncapped 5’ triphosphates and RNAs lacking 2’-O methylation at the first nucleotide (Cap0) [19], as well as short dsRNA regions near the 5’ terminus [27]. TLR3 is an endosomal pattern recognition receptor (PRR), which selectively binds to long dsRNA duplexes (>40nt) [28]. Both receptors signal through the Interferon Responsive (IRF) pathway [29].

IRF-responsive luciferase reporter signaling was measured in THP1 monocytes and two HEK lines stably overexpressing RIG-I or TLR3, following mRNA transfections. THP1 monocytes and HEK-TLR3 cell lines produced the strongest IFN signaling response to unpurified WT T7 mRNA, and a reduced response to HPLC-purified mRNA from the same IVT(Fig 5A). Crude mRNA produced by T7-68 produced even lower responses in these two cell lines, consistent with the prior observation that 3’ extensions and total dsRNA were reduced in the untreated T7-68 sample. HEK-RIG-I cell lines responded with IFN signaling to crude WT-IVT RNA while showing lower responses to mRNA produced by T7-68 or purified WT-produced mRNAs (Fig 5A). Because capping was very high in these samples (99% by LC-MS), the RIG-I response observed is not likely due to differences in uncapped 3’ triphosphate mRNA in the crude WT sample.

**Figure 5.**
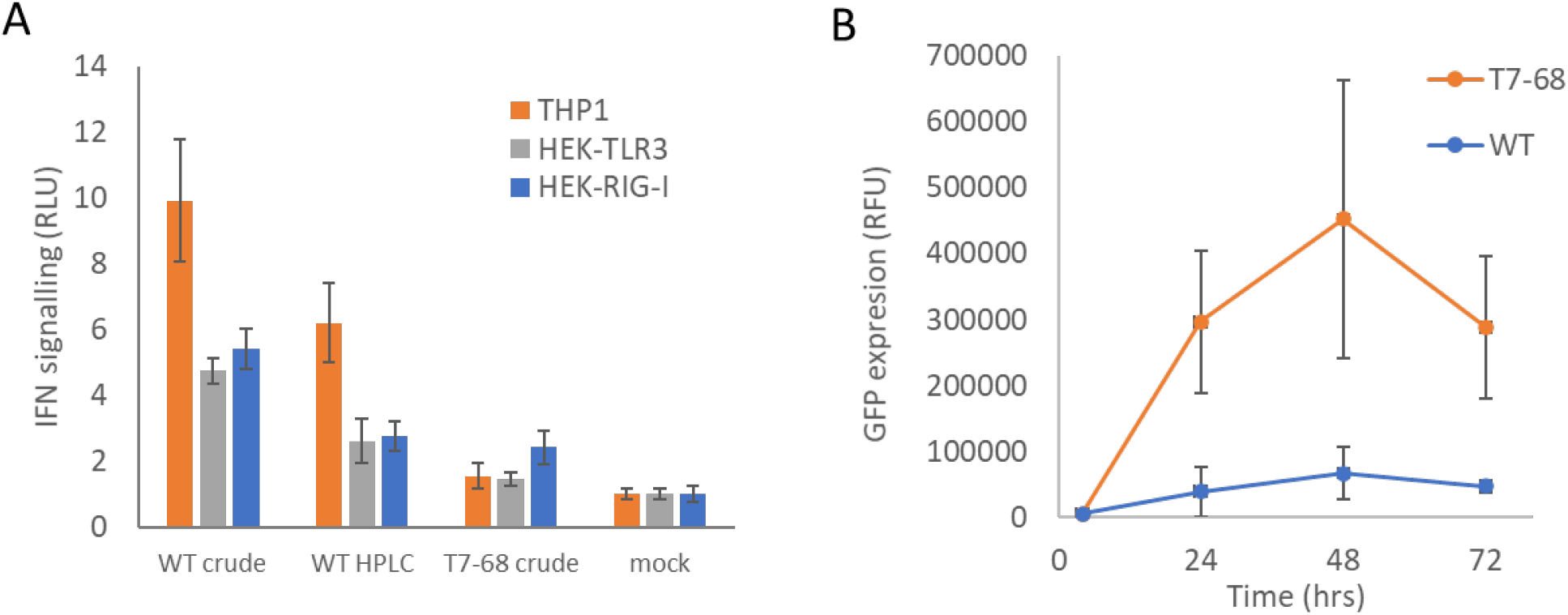
Cell-based reporter assays for immune response and mRNA translation (A) THP1 Dual monocytes, HEK-TLR3, and HEK-RIG-I cell lines with luciferase reporters downstream of an IFN-responsive promoter were LyoVec transfected with mRNA preparations crude and HPLC-purified mRNA preparations analyzed in Figure 5. (B) Crude preparations of eGFP mRNA transcribed using the WT and T7-68 polymerases were transfected into HeLa cells. GFP expression following incubation at was measured by fluorescence emission at 510nm.

Interferon signaling induced by synthetic mRNA inhibits protein translation through PKR phosphorylation of translation initiation factor-2alpha (eIF-2α) [30]. WT T7 and T7-68 polymerases were used to transcribe GFP reporter mRNAs with N1-mψ, and crude samples were used to transfect HeLa cells. Over a 72-hour time course, GFP expression was, on average, 8-fold higher for the crude T7-68 mRNA samples vs the crude WT T7 mRNA sample (Fig 5B). This observation suggests that reduced dsRNA production and IFN signaling in T7-68 mRNAs translated to improvements in protein expression.

## DISCUSSION

The eukaryotic mRNA 5′ cap structure facilitates efficient translation initiation, enhanced mRNA stability, and reduced immunogenicity in vivo [31]. These attributes are critically important to the efficacy and safety profile of mRNA therapeutics, making capping efficiency a critical quality attribute for their manufacture [4]. However, the existing methods for adding native or modified 5’ cap structures enzymatically (via VCE) or co-transcriptionally are both costly and challenging at scale.

We have demonstrated that the evolved T7-68 variant selectively incorporates di- and tri-nucleotide 7-mG cap analogs over GTP at transcription initiation, which allowed for a reduction in cap loading and improved final yields. Because the WT T7RNAP is not selective for dinucleotide cap analogs over GTP at initiation, a ratio of 4:1 cap analog to GTP only achieves ∼70% capping efficiency [5]. The low GTP concentrations required for this condition reduce total mRNA yield. This presents a tradeoff between capping efficiency and mRNA yield, determined by the ratio of Cap:GTP and further limited by the overall cost of the cap. T7-68 was evolved for selective incorporation of the sCap (7mGppp7mG) analog over GTP during transcription initiation. Significant increases in capping were observed at all sCap:GTP ratios tested. T7-68 achieved high capping efficiencies (>95%) in batch IVT reactions, which were not possible with the WT RNA polymerase (∼80%) using both riboswitch and luciferase reporter mRNAs. Importantly, this did not require a tradeoff between increased capping efficiency and yield, as luciferase mRNA yields were equivalent between the WT and T7-68 polymerases at all GTP concentrations tested. T7-68 also demonstrated equivalent yield and capping efficiency when either the sCap or ARCA were used, suggesting that T7-68 has broad substrate recognition for 7mG cap analogs. These dinucleotide cap analogs are established reagents for co-transcriptional capping [17] [16]. Several have been characterized that allow for increased protein expression and half-life in vivo [17] [32] [33]. The benefits of this chemical diversity in synthetic cap analogs are not available when using enzymatic capping, which only produces native Cap-0 mRNA.

T7-68 enabled capping reactions with lower cap inputs for a given desired capping efficiency, with a corresponding increase in mRNA yield in batch reactions. A fed-batch IVT reaction can be used to further increase IVT yield [34] and mRNA capping by maintaining a low steady-state concentration for GTP (∼1mM). This strategy maintains a high ratio of Cap:GTP throughout the reaction, while allowing the reaction to reach completion with GTP in equal stoichiometry to the other nucleotides. The T7-68 polymerase is complementary with this strategy, allowing higher capping efficiencies in fed-batch IVT than would otherwise be achieved with the WT polymerase.

While T7-68 was evolved under selection for incorporation of a dinucleotide cap analog (sCap), we found that T7-68 also incorporates the trinucleotide cap analog, CleanCap AG, with greater efficiency than WT T7. CleanCap AG (m7GpppA_m_G), has several useful features for co-transcriptional capping. It is selectively incorporated over GTP, achieving 94% or higher capping efficiency when used with WT T7 RNAP [18]. The analog also has a 2’-O-methyl modification, yielding a native Cap-1 structure, without the requirement of a separate 2’-O-methyltransferase reaction [5]. CleanCap AG is used for large-scale manufacturing of the BioNTech BNT162b2 SARS-Cov 2 vaccine, demonstrating its scalability and wide utility [35]. We observed ∼97% capping for T7-68 reactions using 1mM CleanCap AG, comparable to the efficiency we observed for 4mM CleanCap AG with the WT T7 RNAP, and consistent with the literature value of 94% [18]. The difference between T7-68 and the WT polymerase were even greater when compared in the same reaction buffer, indicating that the improved capping efficiency is an inherent characteristic of T7-68. A 4-fold reduction in cap loading significantly improves the economics of co-transcriptional capping using CleanCap AG, without sacrificing capping efficiency or yield.

In addition to capping efficiency and final yield, mRNA process development must also consider unwanted byproduct generation. Double-stranded RNA is a byproduct generated by T7 and other phage polymerases as a result of two mechanisms: Short antisense RNAs and longer 3’ extension products resulting from RNA-templated nucleotide addition in cis or trans [24]. Rabideau et al. proposed that mutations in the C-helix or C-linker regions may affect large conformation shifts which occur when the polymerase transitions from the initiation to the elongation state, or change the interaction of the nascent transcript with the polymerase exit tunnel. They demonstrated that S43A and G47A mutations in the C-helix reduced dsRNA production in IVTs [36].

The mutations in T7-68 were directly selected for increased capping efficiency during transcription initiation, and we speculate that these coding mutations significantly remodel the active site. Interestingly, none of these mutations fall inside the C-helix and C-linker regions of the polymerase, but it is possible that they affect the conformational shift to the elongation complex after initiation similarly, thus leading to lower dsRNA formation. With respect to selective incorporation of the cap analog, these changes to the active site may affect the binding of cap analogs to the ternary complex during initiation, elongation of the cap during abortive initiation, or the conformational shift to the elongation state required to enter processive elongation.

Chromatography methods have been used downstream of IVT to remove dsRNA byproducts produced by viral RNA polymerases. Reverse-phase HPLC methods were first used to demonstrate the presence of 3’ extension dsRNA products, and preparative HPLC has been effectively used to remove dsRNA byproducts from small-scale reactions [37]. This method is the gold standard for dsRNA removal and mRNA deimmunization; however, HPLC purification methods do not scale well, limiting their use in production of therapeutic mRNAs [13]. A recently developed method takes advantage of selective binding of dsRNA contaminants to cellulose resin in buffers containing ethanol. Using cellulose chromatography, reductions in residual dsRNA contamination (>90%) and mRNA immunogenicity were equivalent to HPLC purification [14]. While cellulase chromatography scales well for production, recovery rates depended on mRNA length, and were >65%. While effective at removing dsRNA produced by WT T7RNAP, chromatography methods all result in mRNA yield losses, and add complexity and cost to mRNA production.

T7-68 has inherently lower dsRNA byproduct formation under standard IVT conditions. We observed fewer 3’ extension species and a 50-fold reduction in total dsRNA in crude mRNA produced by T7-68, compared to the WT polymerase. Importantly, the 3’ extension products and total dsRNA produced by T7-68 were equivalent to residual contaminants following HPLC-purification of mRNA produced by WT T7RNAP.

dsRNA-mediated immune stimulation was also reduced in crude, unpurified mRNA produced by T7-68. In THP1-Dual, HEK TLR3 and HEK RIG-I cell lines, mRNA produced by WT T7RNAP and HPLC purified generated reduced signaling through an IFN response promoter. Surprisingly, IFN signaling was further reduced 2-fold for the T7-68 crude sample relative to the purified WT mRNA sample, despite similar levels of 3’ extensions and total dsRNA observed by ELISA in these two samples. One possible explanation is that dsRNA produced by T7-68 may be present as shorter antisense duplexes (<40nt), which may be detectable in the sandwich ELISA assay, but do not stimulate TLR3 activation [38].

Consistent with reduced IFN signaling, expression of a GFP mRNA was increased for crude mRNA produced from T7-68 relative to WT. Because these two samples were both 99% capped, this increase in expression is not likely due to differences in cap recognition during translation initiation. It is likely that the reduced IFN signaling from the T7-68 sample improved protein translation by preventing phosphorylation of eIF-2α. Taken together, the reduced dsRNA production, immune stimulation, and translation efficiency observed for crude T7-68 mRNA in this study suggest that the polymerase could simplify mRNA manufacturing by removing the need for chromatography steps, which are otherwise employed after in vitro transcription with WT T7RNAP. This benefit may be used in conjunction with co-transcriptional capping, but could also be employed with enzymatic capping processes using VCE.

An evolved thermostable RNA polymerase (Ts T7-1 or Hi-T7, NEB) has been shown to reduce dsRNA byproducts when used at elevated reaction temperatures. In IVT reactions at 50°C, Ts T7-1 produced fewer small anti-sense and 3’ extension dsRNA contaminants. Moreover, high-temperature IVT products also showed reduced INF-α signaling responses in cell-based assays [15]. However, RNA is prone to hydrolysis at elevated temperatures in the presence divalent cations, including Mg^2+^, commonly used in IVT reactions [39]. Hydrolysis resulting in internal cleavage must be minimized to maintain RNA integrity (the fraction of full-length mRNA in a preparation), a critical quality attribute for therapeutic mRNA manufacturing [40]. Reducing the MgCl_2_ (4mM) and NTP concentrations (0.5mM), as well as the reaction time (1hr) can mitigate hydrolysis of mRNA-length transcripts at 50°C during the IVT reaction, but this approach limits overall mRNA yield. In contrast, we have shown that the T7-68 polymerase produces lower dsRNA byproducts than WT T7RNAP under reaction conditions and temperatures (37°C) that are widely used in mRNA manufacturing and maintain mRNA yield and stability.

We have demonstrated that an engineered RNA polymerase, T7-68, has the potential to simplify mRNA manufacturing through improvements in co-transcriptional capping and reduced dsRNA formation. The increased capping efficiency of T7-68 shifts the existing paradigm and will enable more mRNA manufacturing processes to deploy this simplified, co-transcriptional capping strategy. Protein engineering has the potential to solve outstanding problems in mRNA manufacturing, including yield, processivity, fidelity, and incorporation of novel nucleotides and cap analogs.

## MATERIALS AND METHODS

### In vitro transcription

For polymerase capping characterization, reactions using the ARCA (TriLink) and sCap (Glycosyn) analogs were run in Buffer A (50 mM Tris HCl pH 7.9, 30 mM MgCl_2_, 10 mM DTT) with 5 mM ATP, 5 mM CTP, 5 mM UTP, 1U/μl RNase inhibitor, 0.002 U/μl yeast pyrophosphatase (New England Biolabs), and 50ng/μl linearized DNA template. For cap analog titrations, the total of the cap analog and GTP concentration was 6mM to vary the analog:GTP ratio. Highly purified commercial preparations for the WT T7RNAP (NEB, 12 U/ul final concentration) and T7-68 (Codexis/Alphazyme Codex® HiCap RNA polymerase, 1x final concentrations) were used for polymerase characterization. Reactions ran for 2 hours at 37°C, to ensure high RNA yield and integrity, and were quenched with 60mM EDTA (final).

IVT reactions with CleanCap AG were performed in either Buffer B (30mM Tris pH 8, 27 mM MgCl_2,_ 3 mM DTT) or Buffer C (40mM Tris-HCl pH 8, 10mM DTT, 2mM spermidine, 0.02% (v/v) Triton X-100, 16.5mM Mg Acetate). Buffer B was optimized for yield of T7-68 reactions on long templates up to 6kb, and Buffer C is the buffer recommended by the CleanCap® AG manufacturer (Trilink) for use with WT T7RNAP. The remaining reaction components were 5 mM each nucleotide (ATP, CTP, UTP, GTP), 1U/μl RNase inhibitor (NEB), 0.002 U/μl yeast pyrophosphatase (NEB), 50ng/μl linearized DNA template, and 0-4mM CleanCap AG. The luciferase DNA template for these reactions had the required AGG initiator nucleotides downstream of the promoter to allow incorporation of the CleanCap AG analog.

The mRNA used for 3’ extension, dsRNA characterization, and immunogenicity assays encodes an engineered, site-specific endonuclease based on the I-CreI homing endonuclease from *Chlamydomonas reinhardtii* [41]. IVT with WT T7 RNA polymerase and T7-68 were performed with 5 mM and 2mM CleanCap AG, respectively. IVT reactions were treated with TURBO DNase (ThermoFisher) to digest plasmid DNA. mRNAs were then purified by LiCl precipitation and treated with Phosphatase (New England Biolabs) followed by buffer exchange into 1 mM sodium citrate, pH 6.5. HPLC purification was carried out with Prep scale PLRP-S column (Agilent) using a method adapted from a previous study [37]. HPLC fractions were screened by dsRNA ELISA and HPLC purity assay. Fractions with low dsRNA content and high purity were pooled followed by buffer exchange in 1 mM sodium citrate, pH 6.5. Affinity purification of mRNA was performed using and Oligo dT (18) monolith column (BIA Separations).

Yield experiments for longer 4kb and 6kb IVT templates were performed using T7-68 with buffer B, or the WT T7RNAP with its recommended buffer from the manufacturer supplemented with additional MgCl_2_ to allow for higher yields (NEB, 40 mM Tris-HCl pH 7.9, 30 mM MgCl_2_, 2 mM spermidine, 1 mM dithiothreitol). 5mM N-1-methylpseudouridine was used in the place of UTP. The remaining reaction components were 5 mM each nucleotide (ATP, CTP, GTP), 1U/μl RNasin inhibitor, 0.002 U/μl yeast IPPase (New Englad Biolabs), 50ng/μl linearized DNA template, and 1.5mM CleanCap AG (TriLink).

### Yield assays

IVT yield was assayed using the Quant-iT™ broad range RNA assay kit (Thermo Fisher) according to the manufacturer’s instructions, using a total assay volume of 100μl and standards provided with the kit.

### mRNA capping assays

*In vitro* transcriptions for the capping assay were performed as described previously, using the riboswitch or firefly luciferase templates. Quenched IVT reactions were prepared using the Monarch® RNA cleanup kit 500μg (New England Biolabs), and the concentration of the mRNA eluted in 50μl was determined using the Quant-iT™ broad range RNA assay.

The riboswitch mRNA was autocatalytically cleaved with the addition of ligand. In a 30 μl reaction, 2μM of mRNA was cut in 20mM Tris, pH 7.5, 10mM MgCl2, and 2mM glucosamine-6-phosphate for 30 min at 37°C. The reaction was quenched with the addition of 30μl formamide. The luciferase mRNA was cleaved using a site-specific DNAzyme (5’- CTTCTTTTTCCGAGCCGGACGACTCTTATTT -3’), releasing a 13nt fragment from the 5’ end of the mRNA. In a 24 μl reaction, 2μM mRNA was annealed with 10μM DNAzyme in 5 mM Tris pH 7.5, 15 mM NaCl, 0.1 mM EDTA. The sample was melted at 95°C for 3 min, then snap cooled on ice for 2 min. 3ul each of 10X Cleave buffer (500 mM Tris pH 7.5, 1.5 mM NaCl) and 10X MgCl_2_/MnCl_2_ buffer (100 mM MgCl_2_, 100 mM MnCl_2_) were added to the sample to initiate cleavage. The reaction was incubated at 37°C for 3 hours, then quenched with the addition of 30ul formamide.

Cleaved samples were denatured at 65°C for 5 min, then snap cooled on ice before loading onto a 15% polyacrylamide Urea-PAGE gel. The gel was run for 105min at constant 16W constant power, then stained with 3x GelGreen staining solution (Biotium) for 30-60 minutes in 1x TBE buffer. The gel was imaged using blue light excitation and an ethidium bromide emission filter. Samples were analyzed by gel densitometry using GelAnalyzer 19.1 (www.gelanalyzer.com) by Istvan Lazar Jr., PhD and Istvan Lazar Sr., PhD, CSc.

### Transcription fidelity assay

Polymerase fidelity was measured based on directly sequencing a large number of RT-PCR clones derived from mRNA transcribed from variant polymerases. *In vitro* transcription reactions were performed as described for HTP screening, using 0.5 mM sCap G, 5.5 mM GTP, but omitting glucosamine-6-phosphate. The inclusion of sCap allowed the measurement of the contribution of this cap analog (if any) to the error rate of WT and variant polymerases under process-relevant conditions. A 1.7 kb firefly luciferase DNA template DNA was used with wild-type and variant T7 RNAPs to generate full-length mRNA transcripts. RNA was isolated using the Zymo RNA Clean and concentrator-25 kit (Zymo Research), and residual DNA was removed from the RNA samples by two treatments with the DNA-free DNase I kit (Thermo Fisher), per the manufacturer’s instructions. Samples were reverse-transcribed with Accuprime Reverse Transcriptase (Agilent) using an oligo-(dT)_25_ primer annealing to the (A) tail on the luciferase template. The RT reaction was then amplified using PHUSION^®^ high-fidelity DNA polymerase using HF buffer (New England Biolabs) via PCR to generate a 1675-bp amplicon, using gene-specific primers (Fw 5’- TCTAGAGGCCAGCCTGGCCATAAGGAGATATACATCGGTACTGTTGGTAAAGCCACC-3’, 5- TAATGAGGCCAAACTGGCCACCATCACCATCAGAGCTTGGACTTTCCTCC-3’) annealing to the luciferase coding sequence. Amplified fragments were digested with BglI (New England Biolabs), ligated into a cloning vector, and transformed to generate single clones in *E. coli*.

Individual clones were picked and sequenced using a two-dimensional multiplex barcoding workflow to >20x coverage depth on the Ion Torrent PGM platform (Thermo Fisher). Reads were demultiplexed and mapped against the expected sequence for the 1632-bp region between the amplification primers. Mutations including small insertion, deletions, and single-nucleotide polymorphisms were observed, and the total number of mutations per base of sequenced mRNA-derived clones was calculated. The expected overall rate of mutations per base based on the literature is the sum of errors due to the T7RNAP (∼0.5×10^−4^), Accuscript reverse transcriptase (6×10^−5^) (Agilent, product literature), and Phusion DNA polymerase after 20 cycles of amplification (1.2×10^−5^) (NEB, product literature). Because the error rate from the RNA polymerase was much higher than the RT and PCR steps, the observed error frequency was primarily caused by RNA polymerase misincorporations.

A one-tailed (right-side) binomial test was used to calculate the probability of sampling the observed number of errors (or greater) given number of bases sequenced if the actual error rate in the experiment is equal to the observed overall error rate for T7RNAP-WT in experiment. The fidelity of a given variant was considered indistinguishable from the WT T7RNAP in this assay for p values greater than 0.05.

### HPLC purity assay

Size based RNA separation using ion-pair reverse phase (IP-RP) chromatography was performed on an Thermo Vanquish UHPLC instrument (ThermoFisher Scientific, Waltham, MA). The chromatography was carried out under denaturing conditions at 65°C, using an analytical HPLC column (DNAPac™ RP - 4 μm – 3.0 × 100 mm, ThermoFisher Scientific) containing porous Divinylbenzene (DVB) Polymer beads. Mobile phase consisted of a two-buffer system - Buffer A contained 100mM TEAA, 1mM EDTA in Water and Buffer B contained 100mM TEAA, 1mM EDTA, 25% Acetonitrile in Water (Triethylammonium Acetate TEAA – 1.0M, Sigma-Aldrich, Acetonitrile – HPLC Grade, Fisher Chemical, EDTA – 0.5M, pH 8.0, Thermo Scientific). The column was equilibrated with 40% Buffer B. mRNA was injected onto the column in a 10μL volume, target mRNA load per injection on column was 100ng as measured by absorbance at A260. mRNA was separated over a linear gradient between 40% to 60% Buffer B over 36 minutes at 0.3 ml/min flow rate. This was followed by a column cleaning step at 100% Buffer B for 8 minutes, before switching back to 40% Buffer B, as a column equilibration step for the next injection. RNA eluting from the column was detected on an Vanquish Diode Array Detector (DAD) at 260 nm. RNA species were separated on the column based on size and hydrophobicity, and the size resolved species were recorded as chromatograms by Chromeleon chromatography software (ThermoFisher Scientific).

### dsRNA ELISA

MULTI-ARRAY Standard 96-well plates (Meso Scale Diagnostics, Rockville, Maryland) were coated overnight at 4°C with K1 anti-dsRNA mouse monoclonal antibody (SCICONS, Budapest, Hungary). Plates were blocked using 5% MSD Blocker A (Meso Scale) for 1h, washed, and then incubated with mRNA samples for 90 minutes. Plates were washed again and incubated for 1h with K2 anti-dsRNA (SCICONS). After another wash, captured dsRNA was detected by incubating with sulfo-tagged rat anti-mouse IgM antibody (Thermo Fisher Scientific, Waltham, MA) for 1h before washing and adding MSD GOLD Read Buffer A (Meso Scale) to the plates. Plates were immediately read on an MSD Quickplex SQ 120 instrument and the data was analyzed using MSD Discovery Workbench software.

### Reporter Cell Line Culture

HEK-Lucia RIGI, HEK-Dual TLR3, and THP1-Dual (IRF-responsive) cell lines were purchased from InvivoGen (San Diego, CA). HEK-Lucia RIGI and HEK-Dual TLR3 cells were cultured in Dulbecco’s modified Eagle’s medium (DMEM) (Life Technologies, Waltham, MA) supplemented with 10% heat-inactivated fetal bovine serum (HI-FBS) (Biowest, Riverside, MO), 1U/mL penicillin-streptomycin (Life Technologies), 6mM L-glutamine (Life Technologies), 200μg/mL Zeocin (InvivoGen), 100μg/mL Normocin (InvivoGen), and either 20μg/mL Blasticidin (InvivoGen) for HEK-Lucia RIGI cells, or 200μg/mL Hygromycin B Gold (InvivoGen) for HEK-Dual TLR3 cells. THP1-Dual cells were cultured in RPMI 1640 medium (Life Technologies) supplemented with 10% HI-FBS (Biowest), 1U/mL penicillin-streptomycin (Life Technologies), 4mM L-glutamine (Life Technologies), 25mM HEPES (Life Technologies), 200μg/mL Zeocin (InvivoGen), 100μg/mL Normocin (InvivoGen), and 20μg/mL Blasticidin (InvivoGen).

### Reporter Cell Line Immunogenicity Assays

HP1-Dual cells were plated into a flat bottom 96-well culture plate (Corning Incorporated, Corning, NY) at a density of 100,000 cells/well and phorbol myristate acetate (PMA) (InvivoGen) was added to each well at a final concentration of 10ng/mL. The plate was placed into a 37°C, 5% CO_2_ incubator for 3 days. On the 4^th^ day, the plate was removed from the incubator and the media was replaced twice 1h before transfection. HEK-Lucia RIGI and HEK-Dual TLR3 cells were plated into a flat bottom 96-well culture plate (Corning) at a density of 50,000 cells/well 1h before transfection. All cell lines were transfected with 500ng/well of mRNA using LyoVec (InvivoGen) and placed into an incubator for 24h. Supernatant from each plate was removed and secreted luciferase was detected by adding Quanti-Luc substrate (InvivoGen) and measured on a SpectraMax iD5 plate reader (Molecular Devices, San Jose, CA).

### In-Vitro Potency Assay

Hela cells were cultured in Eagle’s Minimum Essential Medium (EMEM) (American Type Culture Collection, Manassas, VA) supplemented with 10% HI-FBS (Biowest), 1U/mL penicillin-streptomycin (Life Technologies), and 6mM L-glutamine (Life Technologies). Cells were plated at a density of 10,000 cells/well into a 96-well black, clear-bottom plate the day before transfection. RNAs encoding green fluorescent protein (GFP) were complexed with TransIT (Mirus Bio, Madison, WI) according to the manufacturer’s instructions by diluting in EMEM without serum or antibiotics and incubating for 2-5 minutes. Complexed RNA was added to the cells at a dose of 90ng/well and cells were incubated for either 4, 24, 48, or 72 hours. Cells were washed once with PBS, and 100ul of GFP Assay Buffer (Abcam, Cambridge, UK) was added to each well. GFP expression was measured using a SpectraMax iD5 plate reader (Molecular Devices) using excitation/emission wavelengths of 470/510nm.

---

**S1.**
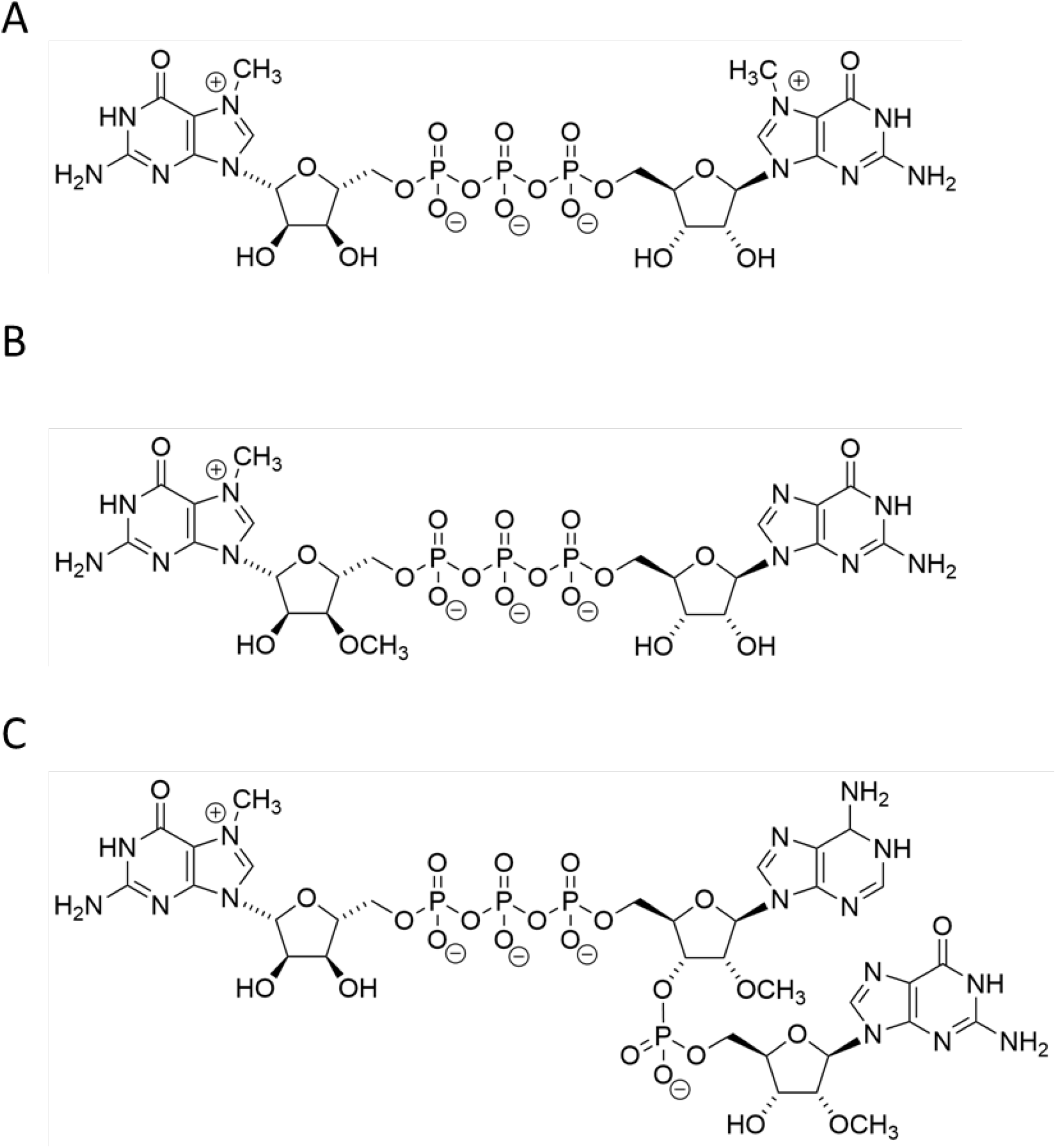
Cap analog structures for (A) sCap, (B) ARCA, and (C) CleanCap™ AG

**S2.**
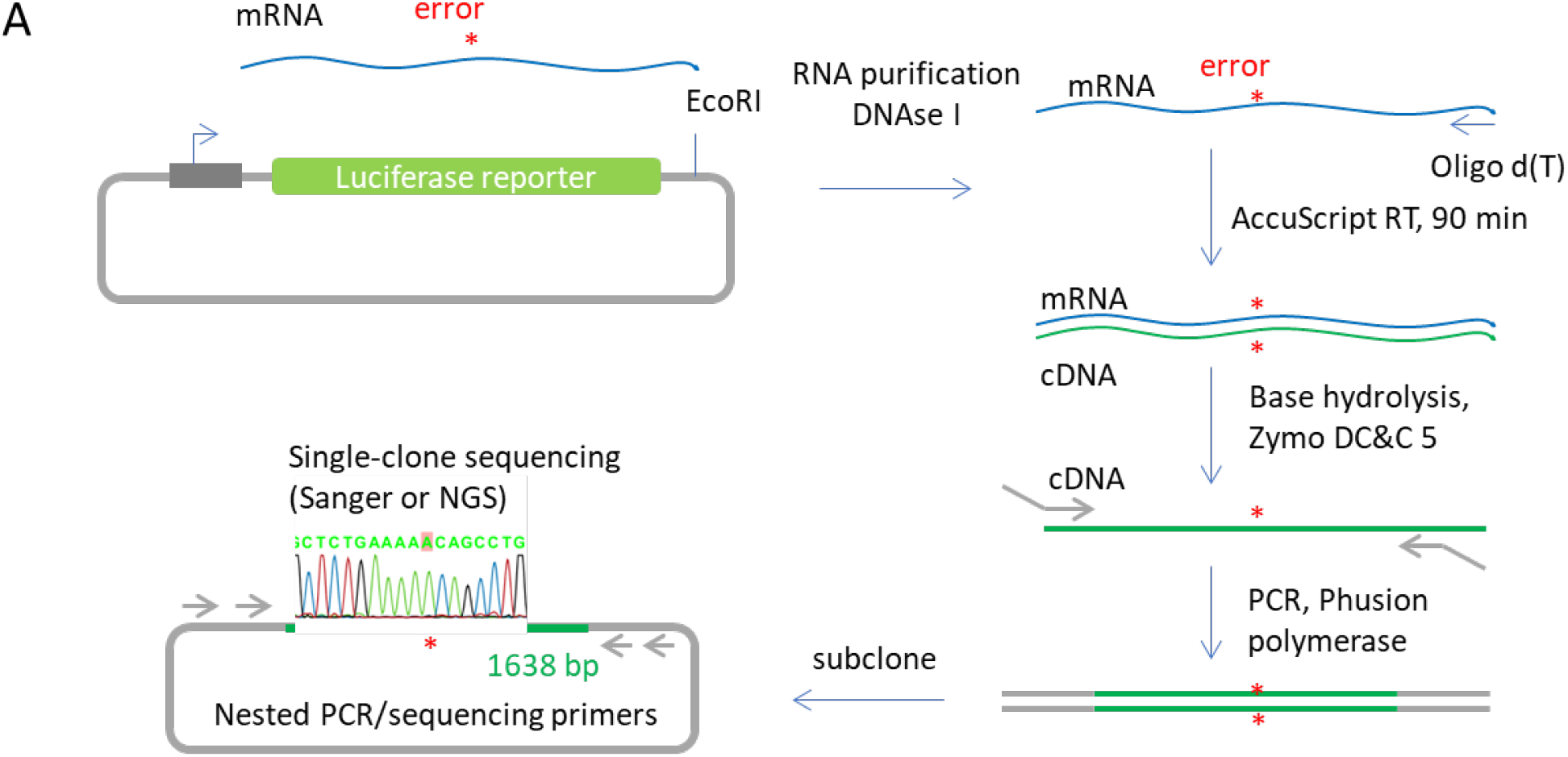
A flow diagram for an RNA polymerase fidelity assay. Polymerase fidelity was measured based on directly sequencing a large number of RT-PCR clones derived from mRNA transcribed from variant polymerases. A 1.7 kb firefly luciferase DNA template DNA was used with wild-type and variant T7 RNAPs to transcribe full-length mRNA transcripts. RNA was isolated and residual DNA was removed from the RNA samples by nuclease treatment. Samples were reverse-transcribed with Accuprime Reverse Transcriptase (Agilent) using an oligo-(dT)_25_ primer annealing to the (A) tail on the luciferase template. The RT reaction was then PCR amplified using PHUSION^®^ high-fidelity DNA polymerase. Individual clones were picked and sequenced to >20x coverage depth on the Ion Torrent PGM platform (Thermo Fisher) in multiplex. Reads were mapped against the expected sequenc, and mutations including small insertion, deletions, and single-nucleotide polymorphisms were observed.

**S3.**
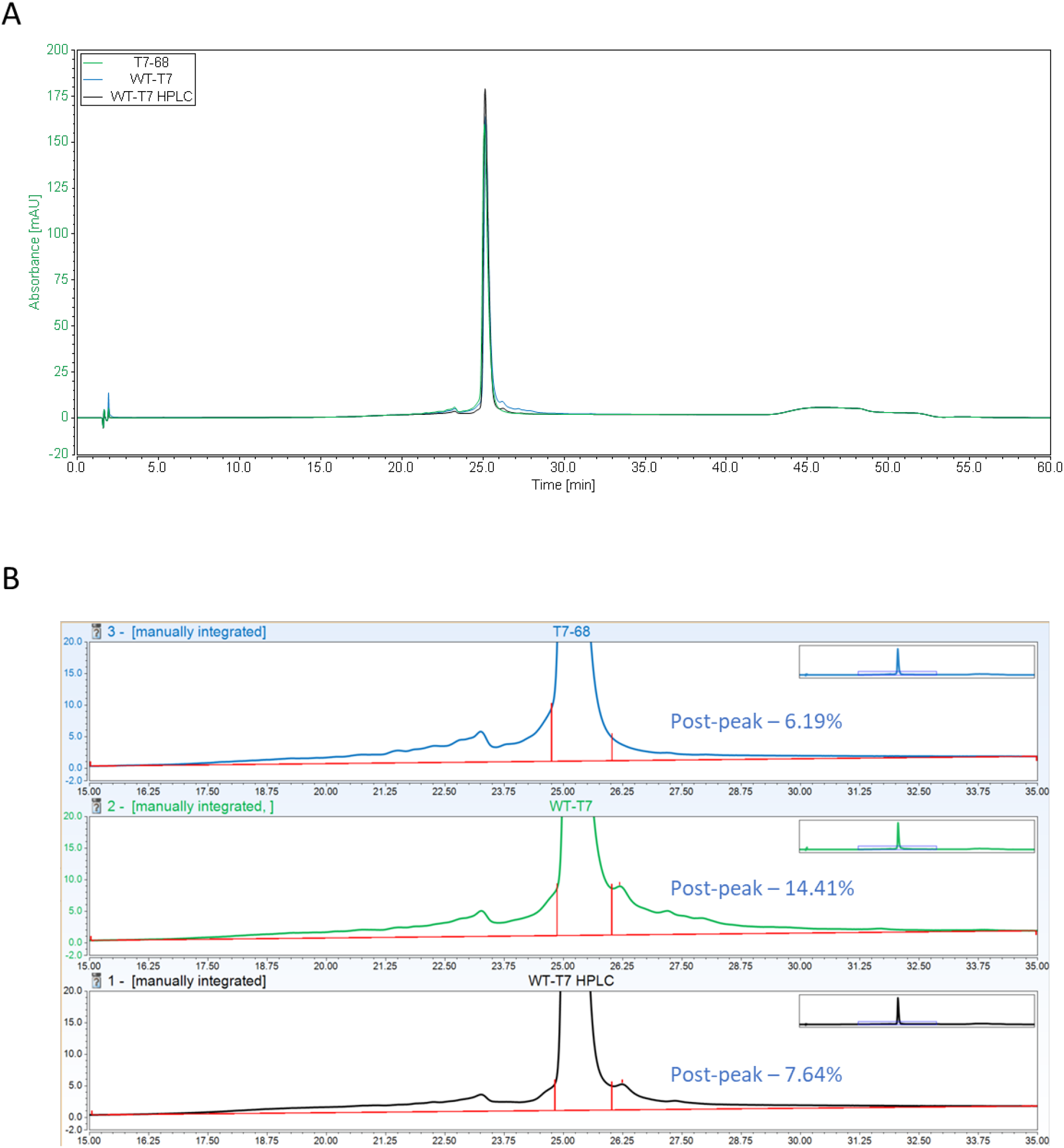
Reverse Phase-HPLC analysis of mRNA 3’ extensions. (A) Full chromatograms for three mRNA samples transcribed using the polymerases indicated (WT, T7-68). The WT sample was analyzed before and after preparative HPLC purification of the mRNA. Equal amounts of each mRNA were injected for each run (B) Integration of apparent higher MW species. Because the baseline does not resolve between the expected mRNA and higher MW species, integrations don’t directly indicate the fractional amount of 3’ extensions in the sample. However, relative comparisons of these numbers between samples can be used to rank their purity with respect to 3’ extension contaminants.

**S4.**
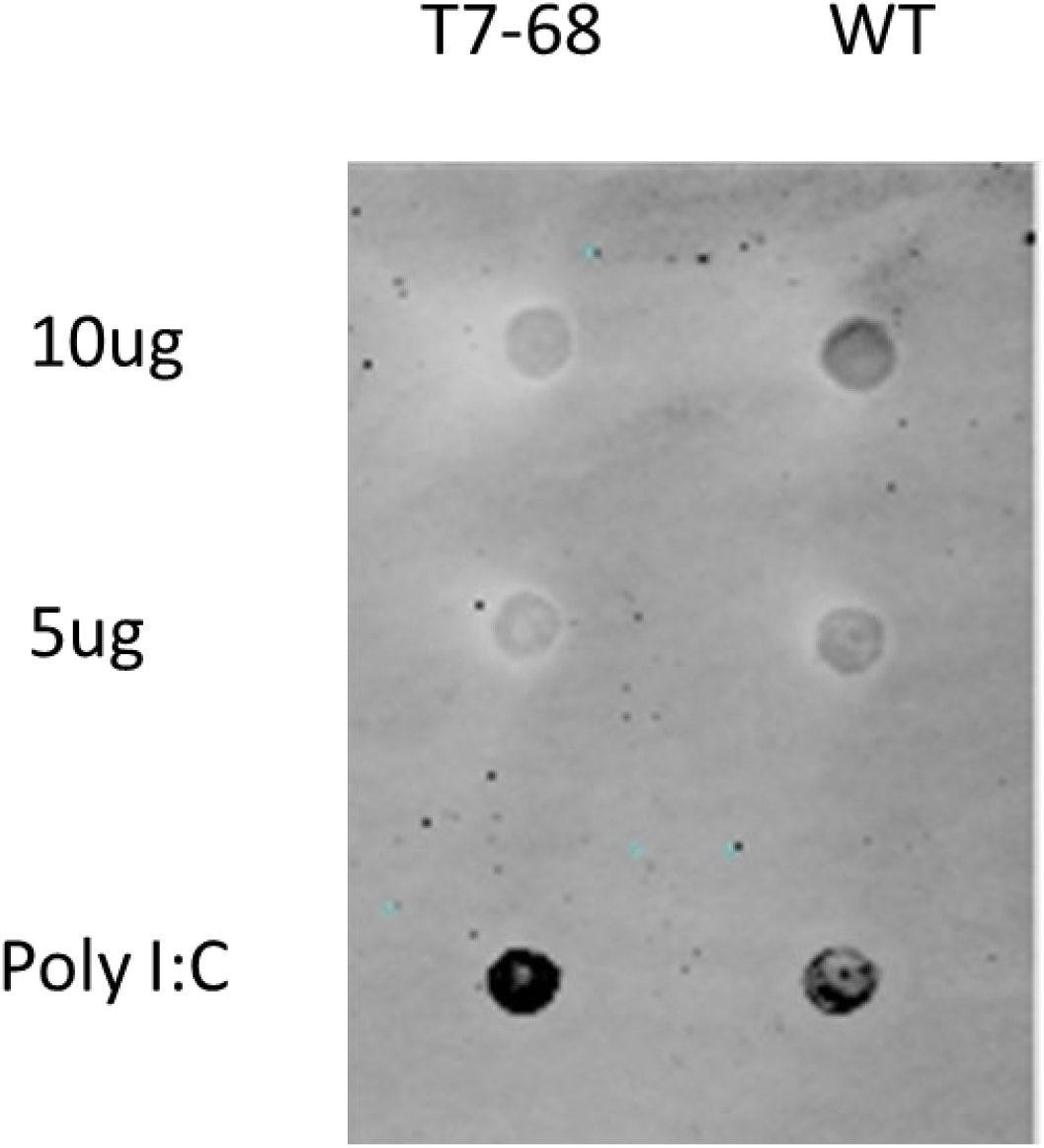
A dsRNA dot blot using the dsRNA-specific J2 antibody. IVT samples transcribed in Buffer B were prepared and spotted onto the membrane in the amounts indicated, and poly I:C was included as a positive control. Higher signal was apparent in crude samples transcribed using the WT RNA polymerase relative to those transcribed using T7-68.

**S5.**
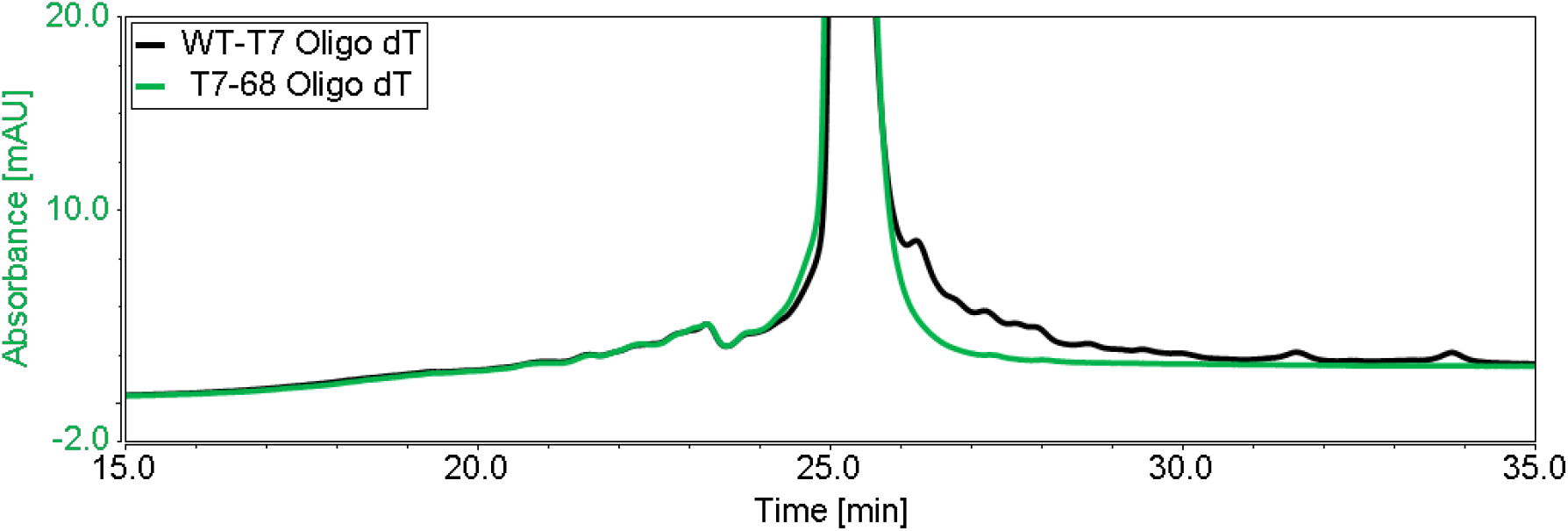
HPLC chromatography of 3’ extension products for WT and T7-68 –produced mRNAs following Oligo-dT affinity purification. Oligo (dT) purification does not remove 3’ extension products, but this scalable process can be used to select for full-length mRNAs. The reduced 3’ extension profile from the T7-68 sample is apparent after Oligo-dT affinity purification.

